# Ribo-macs derived from nucleoli: big ribosome clusters in the cytoplasm of naïve stem cells

**DOI:** 10.1101/2022.04.11.487846

**Authors:** Kezhou Qin, Lei Sun, Xinyi Wu, Jitao Wen, Zhuanzhuan Xing

## Abstract

Primed stem cells and naïve stem cells are important for understanding early development, but their ribosomes have not been focused on. In this study, we find that big ribosome clusters named Ribo-macs exist in the cytoplasm of naïve stem cells. Then, we prove that Ribo-macs are dynamic and physiological in the cytoplasm, and can synthesize proteins associated with biogenesis of endoplasmic reticulum and mitochondria. We also discover and demonstrate that Ribo-macs are the nucleoli, which of significance is promoting us to rethink our understanding of nucleoli. Besides, we reveal that Ribo-macs have a compatible relation with P-bodies and stress granules. In a word, all the results about Ribo-macs provide us with a new insight to understand how cells adapt quickly to environment.

## Introduction

Pluripotent stem cells have two states including naïve state and primed state (*1*), both of which can be switched back and forth by changing the medium (*2*). To identify the molecular factors that control transitions between the two states and establish culture conditions closely recapitulating the signature of human inner cell mass (ICM) cells, researchers have made a lot of efforts in terms of molecular mechanism inducing naïve stem cells from primed stem cells with cytokines or small molecules (*2–8*). In these studies, mutli-omic analysis including DNA Methylomes, transposcriptome, transcriptome, proteome, phosphoproteome, epigenome, metabolome, single-cell transcriptomes (*4, 8–11*) has been applied to explain the whole characteristics of naïve and primed stem cells, but three-dimensional (3D) imaging technology focusing on spatial distribution of ribosomes in cells has not been applied in cell state transition process between naïve state and primed state.

Ribosomes are the central sites for translation in cells and show heterogeneity in controlling selective subsets of mRNAs (*12–14*), and translation control enables cells to quickly respond by directly adjusting protein abundance before a new chromatin state and transcriptional network form, especially dynamic cell fate transitions such as embryonic development (*15*). From these advances, we learned that there are no reports focusing on the relation between ribosomes and other organelle biogenesis such as endoplasmic reticulum (ER) and mitochondria in early embryonic development and naïve stem cells which is important for understanding early development (*2*). In addition, mitochondria with underdeveloped cristae (*16*) and immature ER (*17*) in these stem cells need a large of proteins synthesized by ribosomes to be mature as development and differentiation, so we decided to explore the role of ribosomes using 3D imaging.

Besides, translation control also involves in two other issues including processing-bodies (P-bodies) which mainly play a role in mRNA degradation and translational repression and stress granules consisting of stalled preinitiation complexes that include small (40S), but not large (60S) ribosomal subunits, translation initiation factors eIF4F, eIF3, and PABP, and polyadenylated mRNAs (*18*) in the cytoplasm, both of which can affect translation efficiency. Moreover, stress granules and P-bodies are in a dynamic cycle with actively translating ribosomes and respond rapidly to changes in the polysome-associated pool of ribonucleoproteins (RNPs) that influences the translation and degradation of mRNAs (*19, 20*), which could be a new interpretation why mRNA translation in stress granules is not uncommon (*21*). However, the relation among the three still needs further study.

To our knowledge, there is a high nucleo-cytoplasmic ratio in naïve and primed stem cells. In addition to ribosomes derived from nucleolus, so we also attach great importance to the study of nucleolar region. In the research, we studied the colocalization between ribosomes and mitochondria and between ribosomes and ER using the two methods of immunofluorescence (IF) and the expression of fluorescent protein (FP) fusions and carefully observed the change of nucleolar region based on 3D imaging. In a word, our new findings provide new understanding of ribosomes and nucleoli, and another angle for understanding p-bodies and stress granules.

## Results

### Ribo-macs in the cytoplasm of naïve stem cells

Human embryo stem cells are derived from ICM of blastocysts, induced pluripotent stem cells (iPSCs) are generated through induction of somatic cells, which of both can be maintained in vitro as primed stem cells; there are a few of methods (*2–4, 6, 22–25*) describing how to obtain naïve stem cells, but to avoid the influence of other factors such as feeder and so on, here we choose an easy-to-use, feeder-free and commercial medium called RSeT feeder-free medium to induce and maintain naïve stem cells (Fig. 1A).

**Fig. 1.**
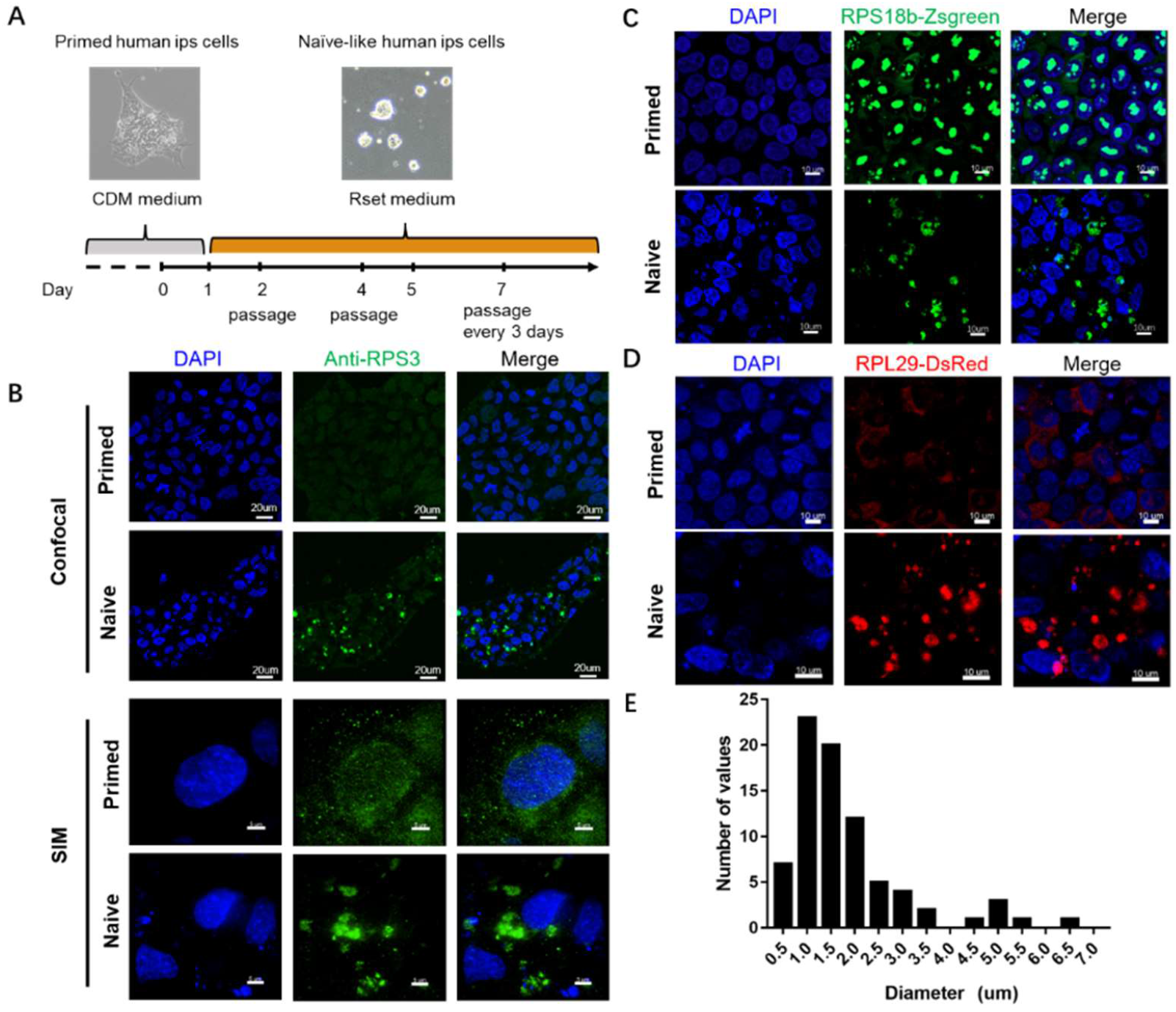
Ribo-macs are firstly discovered in the cytoplasm. (A) Procedure diagram of generation and maintenance of naïve stem cells. (B) Immunofluorescence staining of ribosome small subunit protein RPS3 in naïve and primed stem cells (representative confocal images, scale bar: 20um and representative SIM images, scale bar: 5um). (C) and (D) Representative images in primed and naïve stem cells stably expressing RPS18b-ZsGreen or RPL29–DsRed. Scale bars, 10 μm. (E) Frequency distribution diagram of Ribo-mac diameter.

To determine the role of ribosomes in naïve and primed stem cells, we used two imaging technologies containing IF and FP to visualize ribosome distribution in naïve stem cells derived from an induced pluripotent stem cell line hiPSCs with a primed state. We chose anti-RPS3 antibodies to test endogenous proteins through IF assay and also imaged on super-resolution microscope (SIM) (Fig. 1B, and movie S1 and S2). Meanwhile, we generated a primed stem cell line with overexpression of RPS18b-ZsGreen/RPL29-Dsred, and induce the primed stem cells into naïve stem cells, then imaged on another super-resolution microscope (Zeiss 980) (Fig. 1, C and D). With this analysis, we observed the results of two image-based technologies were the same, that is to say, ribosomes were distributed evenly in the cytoplasm of primed stem cells, but formed big clusters in naïve stem cells which we called macro ribosomes (Ribo-macs).

To further determine the diameter range of Ribo-macs, we plotted the frequency distribution histogram (Fig. 1E); from the histogram, we confirmed the diameter range is between 0.5 um and 7.0 um, and the highest frequency distribution is 1.0 um. Taken together, we find a new phenomenon of ribosome cluster and named the clusters as Ribo-macs.

### Dynamic and physiological Ribo-macs in the cytoplasm

According to the product description of RSeT feeder-free medium, we learned that hiPSCs cultured in RSeT feeder-free medium with a naïve-like state can either be differentiated, or converted back to a primed state. To determine whether the Ribo-macs will disappear when the naïve-like hiPSCs were converted back to the primed state, we plated directly the naïve-like stem cells expressing RPL29-DsRed or not on matrigel-coated plates by culture in CDM medium. After 24 hours, we observed that cell clonal morphology has been changed and the cells looks like primed stem cells (Fig. 2A), then we checked Ribo-macs of these cells using the same imaged-based technologies including IF and FP on super-resolution microscope (SIM). As predicted, we did not observe the Ribo-macs in the cytoplasm of these primed stem cells (Fig. 2, B and C, movie S3 to S6). So we concluded that Ribo-macs will change as the conversion between naïve-like state and primed state.

**Fig. 2.**
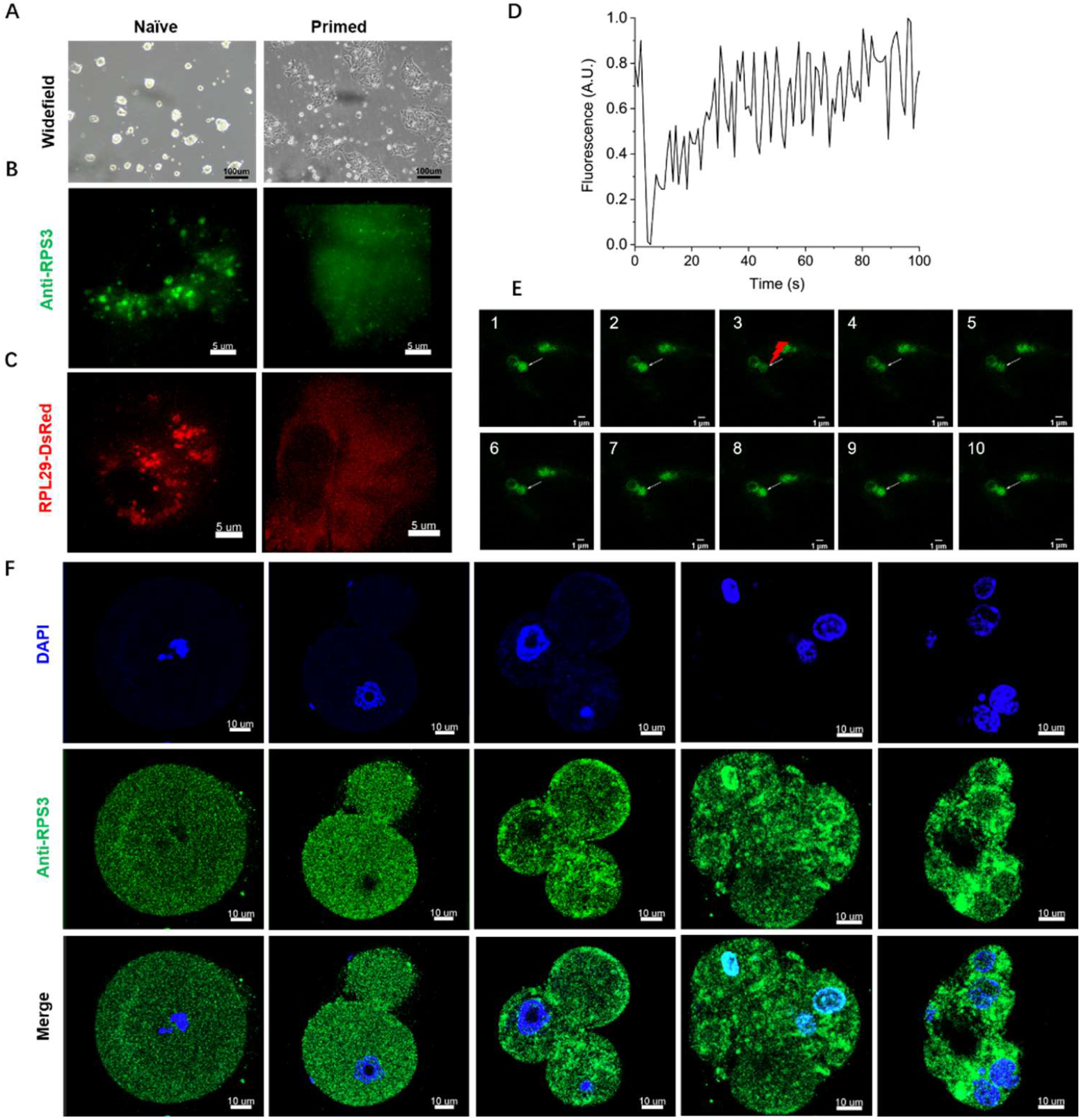
Ribo-macs are dynamic and physiological in the cytoplasm. (A), (B) and (C) Ribo-macs disappeared as naïve stem cells were induced to primed stem cells, which is checked by cell morphology (scale bar, 100um), immunostaining for RPS3 (scale bar, 5um) and expressing RPL29-DsRed (scale bar, 5um). (D) and (E) FRAP of Ribo-macs (white arrowheads, scale bar, 1um). (F) Immunofluorescence staining for RPS3 in 5 stages of mouse embryos, scale bar, 10um.

To further make sure whether the Ribo-macs change dynamically in the cytoplasm of naïve stem cells, we use the fluorescence recovery after photobleaching (FRAP) technology to study protein mobility in the Ribo-macs. We chose naïve-like stem cells with overexpression of RPS18b-ZsGreen to do the FRAP assay and observed that the fluorescence recovered in about 60 second after photobleaching (Fig. 2, D and E, and movie S7). Meanwhile, we learned that the protein BMP4 can reset mouse epiblast stem cells to naïve pluripotency (*26*), so we used 5ng/ml BMP4 to treat primed hiPSCs with expressing RPL29-DsRed for three days and observed Ribo-macs of which fluorescence could be recovered after photobleaching (fig. S1, and movie S8). During the two experiments, we also found the Ribo-macs moved very fast. These results show us the Ribo-macs are dynamic.

To identify whether the Ribo-macs own physiological significance in vivo, we thought that since the naïve-like stem cells are derived from early embryo, the Ribo-macs should exist in the cytoplasm of each stage of embryo development before the blastocyst stage. Then we collected mouse embryos of 5 stages including Zygote, 2 cells, 4-8 cells, Morula, Blastosphere and did immunofluorescence assays using anti-RPL3 antibody (Fig. 2F, and movie S9 to S13). Through comparison analyzation, we found these ribosomes were not distributed evenly in the cytoplasm, and ribosome clusters existed which is similar to Ribo-macs in naïve-like stem cells. So taken together we conclude the Ribo-macs were dynamic and physiological.

### Ribo-macs of rapidly synthesizing proteins

Although protein synthesis by the ribosome can fail in other phase-separation structures such as P-bodies and stress granules, we had a try to check whether the Ribo-macs can synthesize proteins. Firstly, using a stem cell line with co-expression of RPL29-DsRed and RPS18b-ZsGreen, we found the ribosomal large subunit RPL29 and small subunit RPS18b are colocalized together in the Ribo-macs of naïve stem cells (Fig. 3A). Secondly, we followed a published method (*27*) to generate 5 teratomas which have all 3 germ layers including ectoderm, mesoderm, and endoderm confirmed by H&E stains of histology sections and validated that Ribo-macs exist in the 3 germ layers using IF and FP (fig. S2, A to D). We also validated the colocalized phenomenon of ribosomal large subunit and small unit in the Ribo-macs through teratoma 5# (Fig. 3B). Besides, we treated primed hiPSCs with 5ng/ml BMP4 for three days and imaged through IF assay using anti-RPL3 and anti-RPS3. In accordance with the previous results, the large and small ribosomal subunits are also colocalized in the Ribo-macs (Fig. 3C). Taken together, we conclude that the Ribo-macs contain assembled-well ribosomes.

**Fig. 3.**
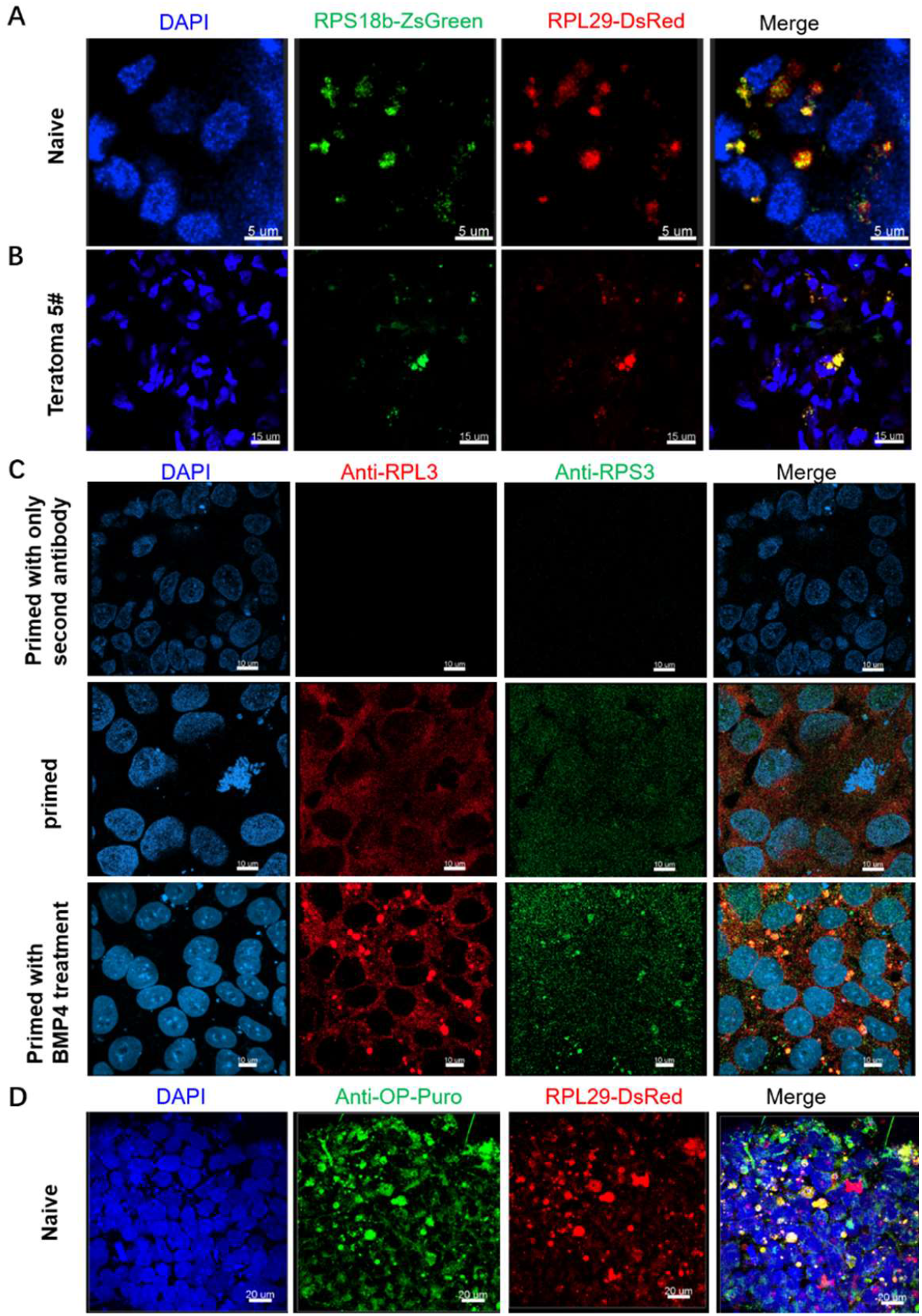
Ribo-macs can produce proteins. (A) and (B) Colocalization of ribosome proteins RPS18b and RPL29 in expressing RPS18b-Zsgreen/RPL29-DsRed naïve stem cells (Scale bar, 5um) and teratoma (Scale bar, 15um). (C) Colocalization of ribosome proteins RPS3 and RPL3 in stem cells treated with BMP4 for three days using IF. Scale bar, 10um. (D) Imaging nascent proteins in expressing RPL29-DsRed naïve stem cells incubated with 30 μM OP-puro. Scale bar, 20um.

To visualize directly the new protein synthesis in the Ribo-macs, we used the alkyne analog of puromycin, O-propargyl-puromycin (OP-puro) to label nascent polypeptide chains (*28*) of naïve stem cells with overexpression of RPL29-DsRed and imaged on super-resolution microscope (Zeiss 980). This result shows us that new synthesized proteins in naïve stem cells incubated with OP-puro for 0.5 hour are colocalized well with the Ribo-macs (Fig. 3D). Overall, proteins can synthesize in the Ribo-macs.

### The relation between Ribo-macs and biogenesis of ER

Since ribosomes are attached to the rough ER, we want to observe the ER morphology and distribution in naïve stem cells. To understand better the relation between Ribo-macs and ER, we generated a naïve stem cell line co-expressing RPL29-DsRed and Sec61b-ZsGreen, and did super-resolution 3D live-cell imaging for 90 min by Zeiss 980. We found that these two markers are colocalized, and the ER signal becomes stronger with the disappearance of ribosome signal, which suggests that the Ribo-macs are synthesizing quickly Sec61b-ZsGreen and gradually depolymerizing (Fig. 4A, and movie S14 to S16). Then we also demonstrated that another ER marker KDEL distributes in the periphery of Ribo-macs on super-resolution microscope (STED) using IF (Fig. 4B), and KDEL-ZsGreen proteins are colocalized with RPL29-DsRed in the cytoplasm of naïve stem cells (Fig. 4C).

**Fig. 4.**
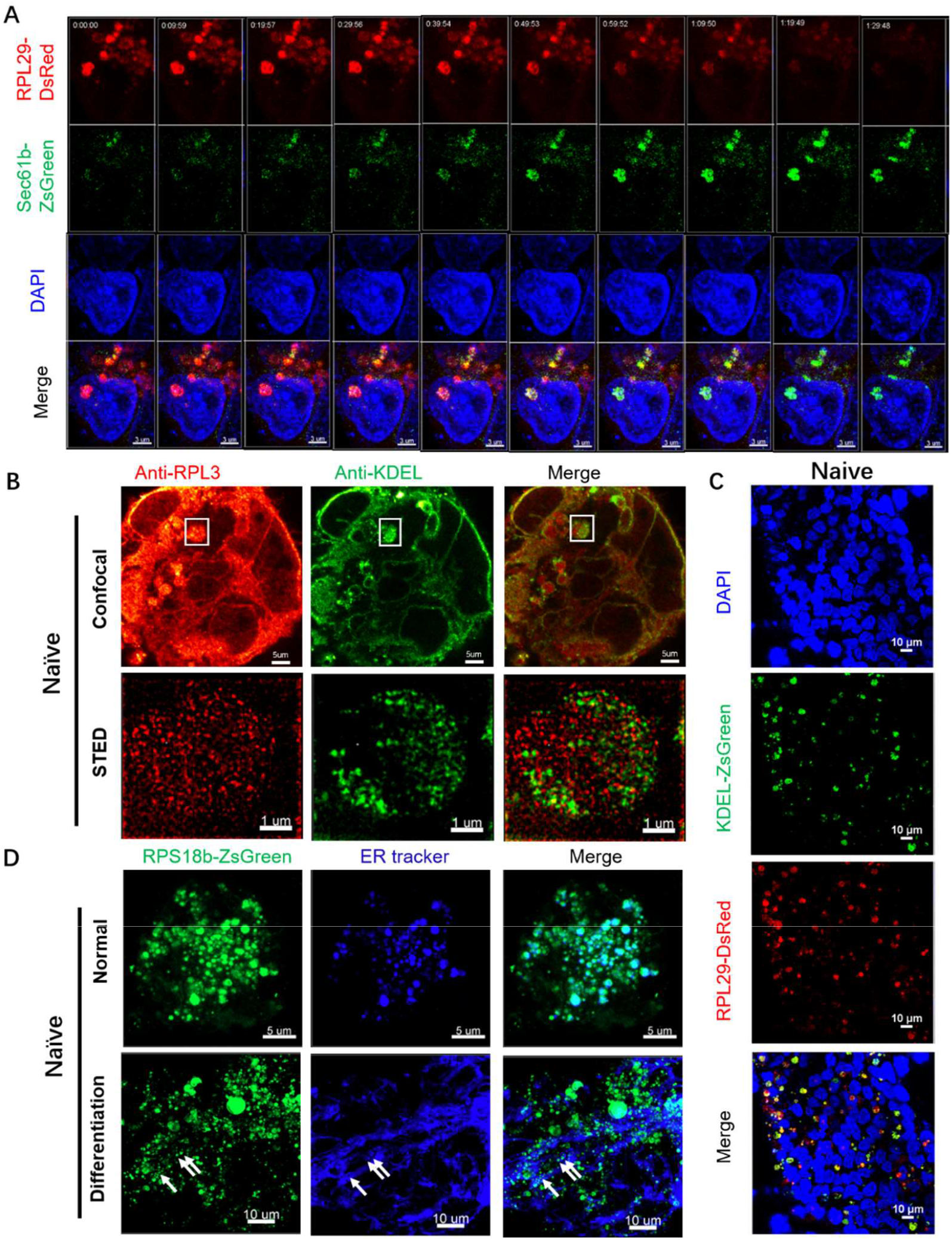
Ribo-macs are related to ER biogenesis. (A) Long-time observation of naïve stem cells with RPL29-DsRed and Sec61b-ZsGreen overexpression, showing that as the expression of protein RPL29-DsRed decreased, the expression of protein Sec61b-Zsgreen increased gradually. Scale bar, 3um. (B) Immunostaining for ribosome subunit protein RPL3 and ER marker KDEL in naïve stem cells and imaging on STED microscope, showing nascent proteins mainly distributed in the edge of Ribo-macs. Scale bar, 5um and 1um. (C) Colocalization of RPL29-DsRed and KDEL-ZsGreen in the cytoplasm of naïve stem cells. Scale bar, 10um. (D) ER staining in expressing RPS18b-Zsgreen naïve stem cells revealed colocalization of RPS18b-ZsGreen and ER, and when naïve stem cells differentiated, Ribo-macs are located in the hole of ER (white arrowheads). Scale bar, 5um and 10um.

To further make sure that Ribo-macs can come out in naïve stem cells and have a relationship with ER, we utilized another kind of naïve medium HENSM (*5*) instead of RSeT medium to induce naïve stem cells expressing RPS18b-ZsGreen, and stained the cells using ER-Tracker dyes. From the imaging, we revealed that Ribo-macs and ER are colocalized completely in naïve stem cells, and reticulated ER is not well colocalized with Ribo-macs in differentiated naïve stem cells but Ribo-macs are located in the hole of ER (Fig. 4D). This result suggested that Ribo-macs play an important role in ER biogenesis.

We also treated primed hiPSCs expressing RPL29-DsRed and Sec61b-ZsGreen with 5ng/ml BMP4 for three days and imaged by STED. As predicted, the Ribo-macs are synthesizing faster the Sec61b-ZsGreen proteins than other ribosomes (fig. S3, and movie S17). Therefore, Ribo-macs are associated closely with ER biogenesis.

### The relation between Ribo-macs and mitochondrial biogenesis

To study the relationship of Ribo-macs and mitochondria, we established a co-expressing RPL29-DsRed and Tomm20-EGFP stem cell line and focused on their colocalization under primed state and naïve state. As expected, we validated the colocalization of Ribo-macs and mitochondria (fig. S4A), which suggests that mitochondrial biogenesis need Ribo-macs. Besides, we also stained an expressing RPL29-DsRed stem cells using Mito-Tracker Green, and imaged at 0 hour and after 3 hours. This result revealed that colocalization exists in between RPL29-DsRed and Mito-Tracker Green at 0 hour, but does not after 3 hours and Mito-Tracker Green clustered to form new mitochondria (fig. S4B).

To further assess the role of Ribo-macs in mitochondrial biogenesis, we selected mouse early developmental embryos as research object, stained the live embryos using Mito-tracker Red and then performed immunofluorescence assay using anti-RPL3. We found Ribo-macs and mitochondria are almost completely co-located in 4 stages (zygote, 2 cells, 4-8 cells, morula) of mouse embryos, and have partial co-location in blastosphere stage suggesting that the mitochondria are maturing (fig. S4C). Taken together, Ribo-macs are associated closely with mitochondrial biogenesis in early embryos.

### The relation between Ribo-macs and the nucleoli that directly come out of the nuclei

As far as we know, ribosome large subunit and small subunit assemble respectively in nucleoli and then come out into the cytoplasm through the nuclear pore. From expressing RPS18b-ZsGreen or RPL29-DsRed primed stem cell lines, we make sure that RPS18b and RPL29 mainly located in nucleoli, but we observed RPS18b and RPL29 mainly located in the cytoplasm of naïve stem cells (Fig. 1, C and D). So we predicted that nucleoli directly come out of the nucleus; thinking about nucleoli are very huge, we did not think the nucleoli come out of the nucleus through the nuclear pore. Here, we mainly focused on two marker proteins including NPM1 which is located in nucleolus and FBL which is located in the dense fibrillar component (DFC) of the nucleolus. To validate the result, we established two stem cell lines, one expressing nucleolus marker NPM1-mCherry (Fig. 5A, and movie S18 and S19) and one co-expressing NPM1-mCherrry and RPS18b-ZsGreen, and compared their localization in primed state and naïve state. As predicted, we visualized the high expression of NPM1 and RPS18b which are colocalized in the nucleoli of primed stem cells and in the cytoplasm of naïve stem cells (Fig. 5B). Besides, we also performed immunofluorescence assay using anti-RPL3 and anti-NPM1 in mouse blastosphere, proved colocalization of RPL3 and NPM1 in the cytoplasm and observed the proteins NPM1 coming out of nuclei (Fig. 5, C and D, and movie S20 and S21), which suggesting nucleoli are coming out of nuclei. Meanwhile, we also selected another nucleolus marker FBL to repeat the immunofluorescence assay in naïve stem cells with expression of RPS18b-ZsGreen, and visualized their colocalization in the cytoplasm or entering to the cytoplasm (Fig. 5, E and F). Taken together, we concluded that Ribo-macs in the cytoplasm are the nucleoli.

**Fig. 5.**
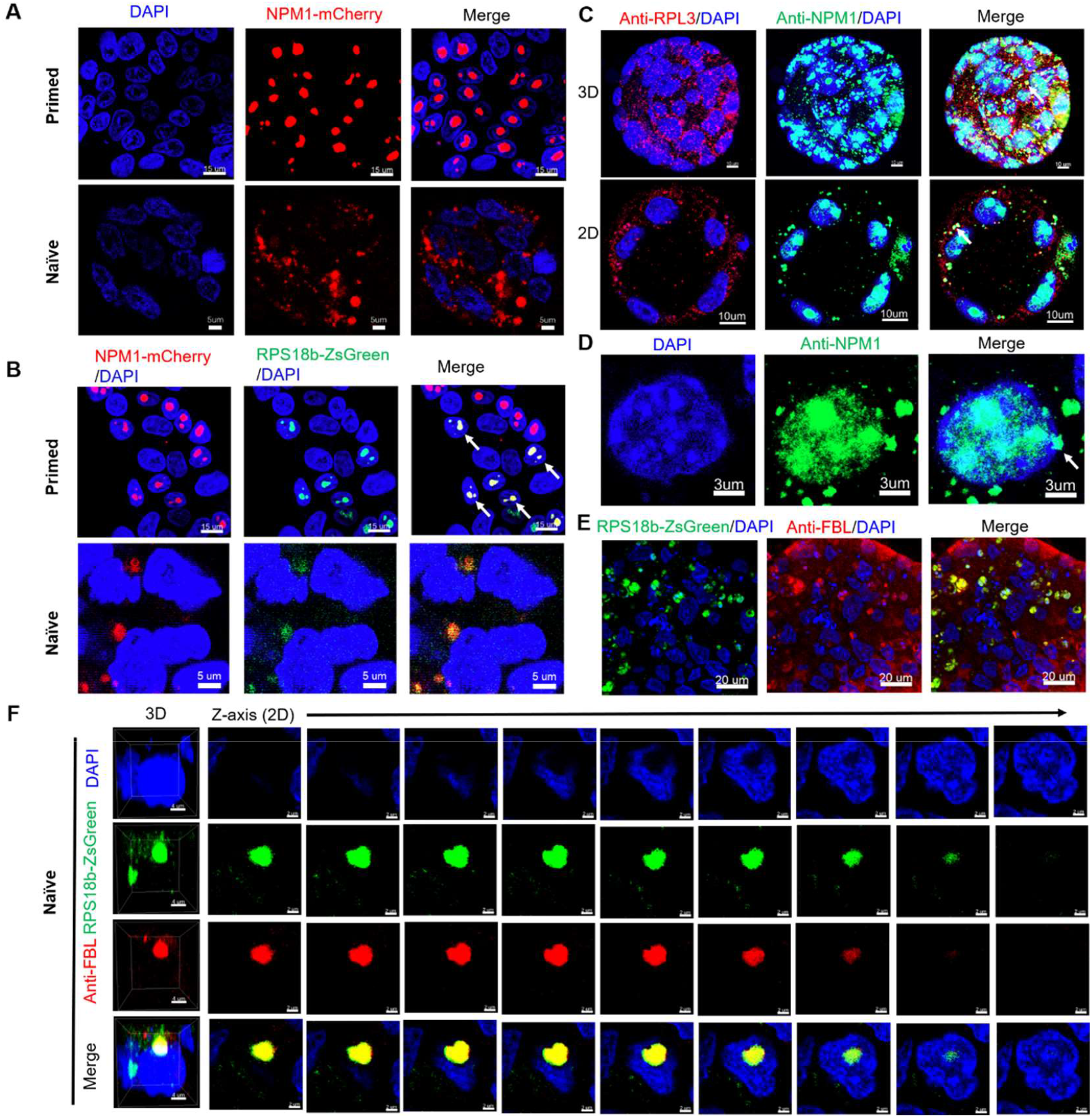
Nuclear export of nucleoli is discovered in naïve stem cells and mouse blastosphere. (A) NPM1 proteins are localized to the nuclei in primed stem cells (scale bar, 15um) and the cytoplasm in naïve stem cells (scale bar, 5um). (B) Colocalization of NPM1-mCherry and RPS18b-ZsGreen in the nuclei of primed stem cells (white arrowheads, scale bar, 15um) and in the cytoplasm of naïve stem cells (scale bar, 5um). (C) Mouse blastosphere were double immunostained for RPL3 and NPM1, and representative images (3D and 2D) show colocalization of RPL3 and NPM1 in the cytoplasm (white arrowheads), scale bar, 10um; (D) Representative images of immunostaining NPM1 in mouse blastosphere showing nuclear export of nucleoli (white arrowhead), scale bar, 3um. (E) Colocalization of FBL and RPS18b-ZsGreen in the cytoplasm of naïve stem cells (scale bar, 20um). (F) Images of FBL and RPS18b-ZsGreen in the direction of Z-axis with 1um interval showing nuclear export of nucleoli, scale bar, 2um.

Since we concluded that Ribo-macs are the nucleoli, we also re-analyzed the other 3D images about RPS18b-ZsGreen, Anti-RPL3 and RPL29-DsRed, and captured several cells in which nucleoli are coming out of the nucleus in naïve stem cells and Teratoma 5# (fig. S5, A to D). Overall, we demonstrated that Ribo-macs are the nucleoli and the nucleoli going out to the cytoplasm, do not depend on the nuclear pore.

### The relation between Ribo-macs and p-bodies and stress granules

Both of p-bodies and stress granules belong to dynamic non-membrane granules in the cytoplasm, so we decided to explore whether they are corelated to Ribo-macs. We firstly generated a LSM14a-EGFP stem cell line to analyze the characteristics of p-bodies in primed state and naïve state. Then, At the time of imaging, ribosomal protein RPL3 was labeled through IF. After analyzing the images, we discovered that Ribo-macs have colocalization with p-bodies in the cytoplasm of naïve stem cells, but to be precise, Ribo-macs contain p-bodies (Fig. 6A). Combined with a RPL29-DsRed stem cell line and stress granule marker antibody anti-G3BP1, we assess the relationship between stress granules and Ribo-macs. The images told us that stress granules and Ribo-macs are colocalized and the state coming out of the nuclei is captured (Fig. 6, B and C). Taken together, we demonstrated that Ribo-macs are related to p-bodies and stress granules, and we speculated that an inclusive relationship between them may be needed for rapid mutual transformation.

**Fig.6.**
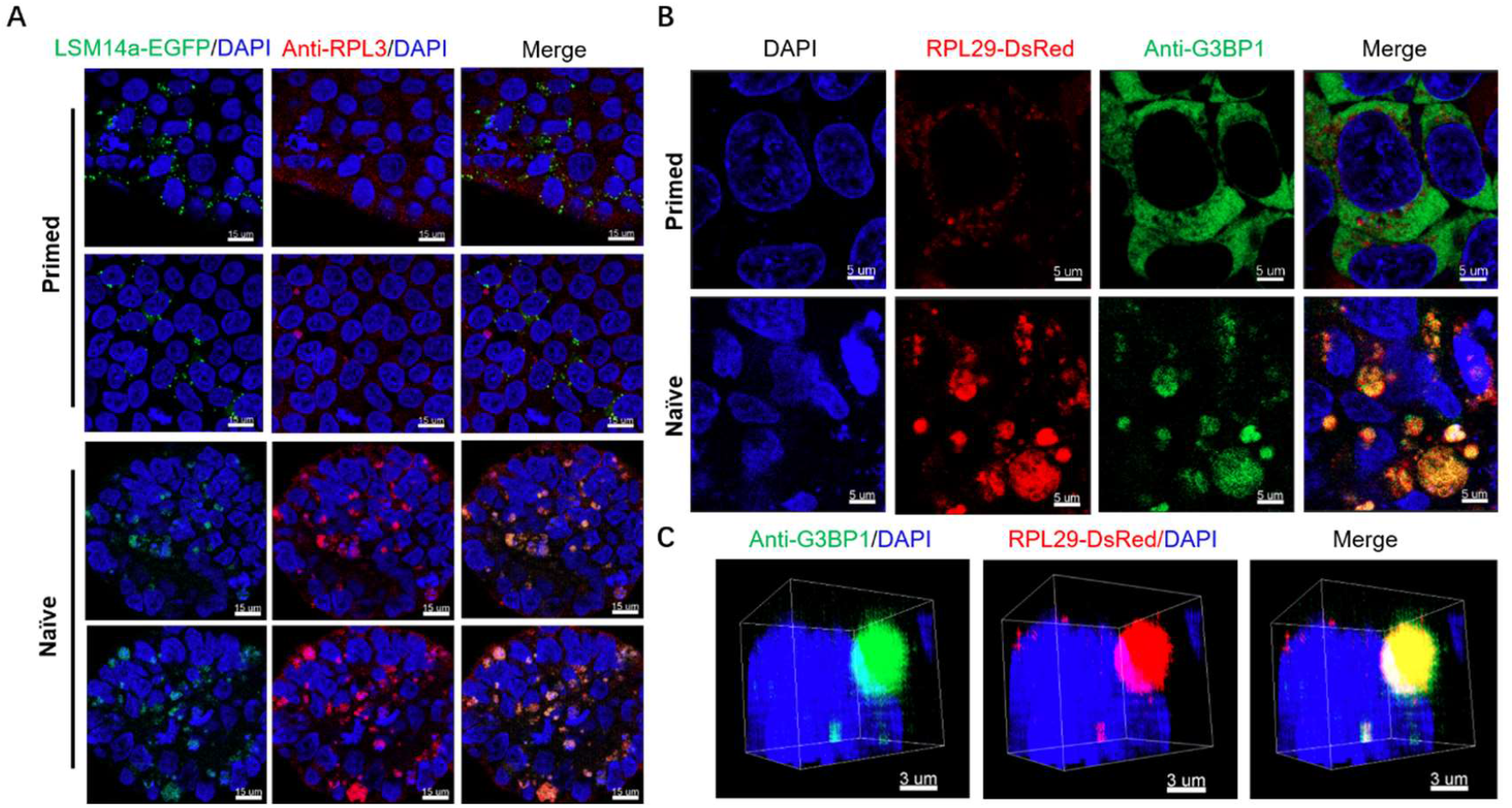
There is a colocalization between Ribo-macs and P-bodies and stress granules. (A) Colocalization of LSM14a-EGFP and RPL3 in the cytoplasm of naïve stem cells but not in the primed stem cells, scale bar, 15um. (B) Colocalization of G3BP1 and RPL29-DsRed in the cytoplasm of naïve stem cells but not in the primed stem cells, scale bar, 5um. (C) A 3D image shows colocalization of G3BP1 and RPL29-DsRed in the cytoplasm of naïve stem cells, scale bar, 3um.

## Discussion

Ribo-macs, which are first discovered and proved to be the nucleoli of nuclei, have an important physiological significance. An article (*29*) reported that nucleoli are removed from the nucleus at the period of zygote, and will be reformed when the mouse embryos develop to 4-cell stage, suggesting that absence of nucleoli cannot result in cell death. Besides, to our knowledge, there are several nucleoli in a nucleus, which of problem has not been focused on. We think that when the nucleoli leave nucleus in response to different stress stimuli, lots of ribosomes also come out together which facilitates the synthesis of large amounts of proteins. We also predict that different nucleoli standing for heterogeneous ribosomes (*12*) may be mixed with different mRNAs in a nucleus, and when they come out of nucleus together for synthesizing lots of proteins needed, it is great for the cells to adapt quickly to the environment. The discovery and proof of nuclear export of nucleoli reminds us that when multiple nucleoli exist in a nucleus, are they the same in composition and function? Meanwhile we believe that the nucleus has a highly regular, highly organized environment, and it is in the cytoplasm that the organized particles from nucleus should perform their proper functions.

## Acknowledgments

We thank professor Yan Qin, professor Jianjun Luo and associate professor Si Wu for support and guide, professor Tao Xu for advice and discussions, Ms Jing sun (the laboratory animal research center) for teratoma formation assay, Junfeng Hao (Core Facility for Protein Research) for tissue section identification at the Institute of Biophysics, Chinese Academy of Sciences. We also would like to thank the Center for Biological Imaging (CBI, http://cbi.ibp.ac.cn), Institute of Biophysics, Chinese Academy of Sciences for help in taking and analyzing images, and the staff from the laboratory animal research center at the Institute of Biophysics for technical assistance and efficient animal care.

## Author contributions

K.Z.Q., X.Y.W., J.T.W. wrote the manuscript. K.Z.Q and L.S. cloned constructs. K.Z.Q and L.S. packaged lentiviruses. K.Z.Q processed all images. K.Z.Q, X.Y.W. and Z.Z.X. performed immunofluorescence assays of mouse embryos. K.Z.Q and J.T.W. performed FRAP assays. All authors discussed the results and commented on the manuscript. K.Z.Q. directed and supervised the project.

## Supplementary Materials

Materials and Methods

Figs. S1 to S5

Movies S1 to S21

