## Supplementary materials for "Ribo-macs derived from nucleoli: big ribosome clusters in the cytoplasm of naïve stem cells"

### Materials and Methods

#### 1. Materials

| REAGENT or RESOURCE | SOURCE | IDENTIFIER |
|---|---|---|
| PD0325901 | GLPBIO | GC10397 |
| human LIF | PeproTech | 300-05-25UG |
| Go 6983 | GLPBIO | GC16907 |
| XAV939 | Sigma-Aldrich | X3004-5MG |
| CGP77675 | Sigma-Aldrich | SML0314-5MG |
| L-Ascorbic acid 2-phosphate sesquimagnesium salt hydrate | Sigma-Aldrich | A8960-5G |
| Dimethyl 2-oxoglutarate | Sigma-Aldrich | 349631-5G |
| Normocin™ | InvivoGen | ant-nr-1 |
| Anti-KDEL antibody [EPR12668] - ER Marker | Abcam | ab176333-10ul |
| RPL3 Mouse Monoclonal Antibody | Proteintech | 66130-1-Ig |
| G3BP1 Polyclonal antibody | Proteintech | 13057-2-AP |
| FBL Polyclonal antibody | Proteintech | 16021-1-AP |
| LSM14A Polyclonal antibody | Proteintech | 18336-1-AP |
| B23/NPM1 Monoclonal antibody | Proteintech | 60096-1-Ig |

| | | |
|---|---|---|
| Anti-RPS3 antibody [EPR7808] - Ribosome Marker | Abcam | ab128995 |
| rRNA (Y10b) | SANTA CRUZ | sc-33678 |
| 35mm glass bottom dish with grid inside, grid size 0.6*0.6mm | Cellvis | D35-14-1.5GI |
| Y-27632 (ROCK inhibitor) | Beyotime | SC0326-10mM |
| Anti-Lamin B1 antibody [EPR8985(B)] | Abcam | ab133741 |
| DH5α Chemically Competent cell | GenStar | S101-02 |
| Corning® Matrigel® Basement Membrane Matrix, *LDEV-Free, 5mL | BioCoat | 356234 |
| Hyaluronidase from bovine testes (Type I-S, 400-1000units/mg solid) | Beyotime | ST1384-25mg |
| RSeT Feeder-Free Medium | STEMCELL | 05975 |
| Lipofectamine 3000 Transfection Reagent | Thermo Fisher | L3000015 |
| Gentle Cell Dissociation Reagent | STEMCELL | 7174 |
| TRYPLE EXPRESS | Thermo Fisher | 12604021 |
| N2 | Thermo Fisher | 17502048 |
| B27 | Thermo Fisher | 17504044 |
| TRYPSIN 0.05% EDTA | Thermo Fisher | 25300062 |
| STEMPRO ACCUTASE | Thermo Fisher | A1110501 |
| hPSC-CDM | Cauliscell Inc | 400105 |
| hiPSCs | Given by Yue Ma lab | Given by Yue Ma lab |
| MEM Non-Essential Amino Acids Solution (100X,100ml) | Gibco | 11140-050 |
| GLUTAMAX I, 100X,100ml | Thermo Fisher | 35050061 |
| NEUROBASAL MED SFM | Thermo Fisher | 21103049 |
| Polybrene (10mg/ml) | Solarbio | H8761 |
| FITC-labeled Goat Anti-Rabbit IgG (H+L) | Beyotime | A0562 |
| FITC-labeled Goat Anti-Mouse IgG (H+L) | Beyotime | A0568 |
| Alexa Fluor 555-labeled Donkey Anti-Rabbit IgG(H+L) | Beyotime | A0453 |
| Alexa Fluor 555-labeled Donkey Anti-Mouse IgG(H+L) | Beyotime | A0460 |
| STAR RED goat anti-rabbit IgG | Abberior | STRED-1002-20ug |
| STAR RED goat anti-rabbit IgG | Abberior | STRED-1001-20ug |
| NucBlue™ Live ReadyProbes™ reagent | Thermal Fisher | R37605 |

| | | |
|---|---|---|
| Mito-Tracker Green | Thermal Fisher | M7514 |
| Mito-Tracker Red | Thermal Fisher | M22425 |
| ER-Tracker™ Blue-White DPX | Thermal Fisher | E12353 |

### 2. Methods

#### 2.1 Cell Maintenance

hiPSCs (given by Yue Ma lab, Institute of Biophysics in Chinese Academy of Sciences) were cultured in hPSC-CDM™(Cauliscell Inc. #400105) supplemented with hPSC-CDM™ supplement (Cauliscell Inc. #600301) or mTeSR™1 Complete Kit (Catalog #85850) on Matrigel-coated (Corning® Matrigel® Basement Membrane Matrix, *LDEV-Free, #356234) 6-well plates and were passaged with 500μM EDTA for 3-5min.

Naïve stem cells were generated, derived and stabilized by RSeT Feeder-Free Medium (Stem cell, #05975) according to user's guide or HENSM (Human Enhanced Naïve Stem Cell Medium) which is devised by Bayerl, J. et al.

#### 2.2 Animals

Animals used in this study were female NOD-SCID mice 4 weeks of age for teratoma formation, female and male C57BL/6J mice with 4-8 weeks for obtaining developmental embryos. The two kinds of animal experiments were performed in compliance with the applicable institutional and/or national guidelines for the care and use of laboratory animals and were approved by the institutional biomedical research ethics committee of the Institute of Biophysics, Chinese Academy of Sciences (permit number: SYXK202108). Mice were group housed (up to 5 animals per cage) on a 12:12 hour light-dark cycle, with free access to food and water in a pathogen-free facility. All mice used were healthy and were not involved in any previous procedures nor drug treatment unless indicated otherwise.

#### 2.3 Conversion of cell state between NV cells and ST cells

According to the instructions of RSeT™ feeder-free medium, primed hiPSCs maintained in RSeT™ Feeder-Free medium were reverted to a Naïve-Like State. On day 1, Primed hiPSCs were plated as aggregates in CDM medium and were cultured at 37°C and 5% CO2 in normal conditions. On day 2, the cells were cultured in hypoxic conditions, the medium was RSeT™ Feeder-Free medium and exchanged every other day. By day 4, the colonies were passaged at a ratio of 1:3 and the cells were passaged every three days. Although during the initial culture in RSeT™ Feeder-Free medium, colonies begin to adopt a highly domed morphology characteristic of naïve-like stem cells, the cells with more than passage 3 were applied to do experiments except special description. When the cells were passaged and cultured in CDM medium at 37°C and 5% CO2, Naïve-like stem cells can be changed into the primed stem cell state, which only need for one day.

**2.4 Immunofluorescence Assay**

hiPSCs and naïve stem cells were fixed with 4% (w/v) paraformaldehyde for 15 min and permeabilized with 0.1% (v/v) Triton X-100 in PBS for 15 min. After blocking with 1% (v/v) BSA (Sangon Biotech, #9048-46-8) and 0.1% Tween 20 (Sinopharm Chemical Reagent Co., Ltd., #30189328) for 60 min, cells were incubated with primary antibodies overnight at 4°C. Cells were washed with PBS for three times, then incubated with secondary antibodies and visualized by super-resolution microscopy including Zeiss LSM980, SIM/GI-SIM, STED and confocal microscopy (Olympus 1000/3000 and Nikon). The images were taken at room temperature and images were analyzed using ZEN 2.6 (blue edition), Imaris and NIS_viewer.

**2.5 Vector Construction**

Total RNA from human ESCs was extracted with Trizol (Life Technologies). RNA yield was determined using the NanoDrop ND-1000 spectrophotometer (NanoDrop Technologies). 1 μg RNA was synthesized to cDNA using the PrimeScript$^{TM}$ RT reagent Kit with gDNA Eraser (TAKARA). Individual genes such as *rpl29*, *rps18b*, *sec61b*, *kdel* and *tomm20* were PCR amplified utilizing overlapping forward and reverse primers custom designed with flanking sequences compatible with the BamHI and EcoRI restriction sites. The lentiviral plasmid pKD-EF1a (an EF1-alpha promoter, ZsGreen or DsRed, IRES domain, and puromycin-resistance gene backbone) for the barcode vector was constructed containing ZsGreen or DsRed transgene flanked by BamHI and EcoRI restriction sites, followed by puromycin resistance enzyme gene downstream. The clonal genes were digested with BamHI-HF and EcoRI-HF (New England Biolabs) at 37 °C for 30 minutes in a reaction system (including 1ug plasmids, Buffer NEB 3.1, 3ul, BamHI-HF, 0.5ul, EcoRI-HF, 0.5ul, H2O up to 50ml), and a 20 bp DNA fragment with flanking sequences compatible with the site of pKD-EF1a vector, was inserted immediately into linearized pKD-EF1α vector by Gibson assembly (Seamless Cloning Kit, D7010M). After screening, the vector was purified using a QIAquick PCR Purification Kit (QIAGEN).

**2.6 Lentiviral Production and Transduction**

According to detailed protocol for lentiviral production using Lipofectamine 3000 reagent (Life Technologies), HEK293T cells were maintained in high glucose DMEM supplemented with 10% fetal bovine serum (FBS). About ~7.2 x $10^6$ cells were seeded in a 10 cm dish in high glucose DMEM with 5% FBS 1 day prior to transfection, such that they were 95%–99% confluent at the time of transfection. For each 10 cm dish, 41ul of Lipofectamine 3000 was added to 1.5 mL of Opti-MEM (Life Technologies) as A. Separately 4.5 ug of pMD2.G, 9 ug of psPAX2, 4.5 ug of an individual vector such as pKD-Flag-IRES2-puro and 35ul P3000 was added to 1.5 mL of Opti-MEM as B. After 5 minutes of incubation at room temperature, the Lipofectamine 3000 (A) and DNA solutions (B) were mixed and incubated at room temperature for 15 minutes. After the incubation period, medium in each dish was removed in half and the mixture (A and B) was added dropwise to each dish of HEK293T cells. Cell supernatant containing the viral particles was harvested after 24 and 52 hours posttransfection, filtered with 0.45um filters (Steriflip, Millipore), and further concentrated using Amicon Ultra-15 centrifugal ultrafilters with a 100,000 NMWL cutoff (Millipore) to a final volume of about 800ul, divided into aliquots and frozen at -80°C.

For viral transduction, virus was added to stem cells at 40% confluency alongside polybrene (5 mg/ml, Millipore) in fresh hPSC-CDM medium. The following day, medium was replaced with fresh hPSC-CDM medium. Selection reagent (puromycin at 0.75mg/mL) was added 48 hours after transduction and was replaced daily for enough ZsGreen or DsRed positive cells.

**2.7 Flow Cytometry**

hiPSCs were transfected using Lentivirus, then ZsGreen or DsRed positive cells were screened, expanded, digested into single cells using accutase (Thermo Fisher, A1110501) and processed using flow cytometry to determine two-positive cells or ZsGreen positive cells or DsRed positive cells.

**2.8 long-time observation of living cells**

Live cells were cultured in confocal dishes, stained by NucBlue™ Live ReadyProbes™ reagent (Thermal Fisher, R37605) for 15min, then washed three times with 1x PBS, and live-cell imaging using live cell Imaging System is mainly performed with super-resolution microscopy Zeiss LSM980 providing fast acquisition. 3D imaging was captured with 1um interval on Z-stack, and long-time imaging was set with 10 min interval to visualize cell dynamics for total 90 min. Images and videos were produced using Imaris soft.

**2.9 Collection of mice embryos at different stages**

Using 12-week-old male mice and 4-week-old female mice (SiPeiFu Biotech Co., Ltd.) of the C57BL/6 strain as donors, PMSG (Phenylmethylsulfonyl fluoride, Ningbo Sansheng, CasNo:329-98-6) 10 IU/mouse was injected intraperitoneally on the first day to promote the maturation of female mouse oocytes. 48 hours later, 10 IU/mouse of HCG (Human Chorionic Gonadotropin, Ningbo Sansheng, CasNo:9002-61-3) was injected intraperitoneally to promote mature oocytes discharged. After 14 to 16 hours, the fresh sperm from the epididymis of the male mouse and the oocytes in the oviduct of the female mouse were taken out, mixed and placed in the pre-heated HTF (Cosmo Bio, cat.no: CSR-R-B070) droplets, which was then placed in a CO2 incubator at 37°C and 5% CO2 for in vitro fertilization. After 4 hours, the embryos with the second polar body as fertilized eggs were washed with M2 (Yiweidi cat.no: M004C), selected, collected and cultured under the same conditions. Fertilized eggs were obtained by in vitro fertilization, and then the embryos of the corresponding period were obtained at differently developmental phase of embryos. After 24 hours, the embryos with round shape and symmetrical division were selected and collected as 2-cell. After 48 hours, the embryos with round shape, symmetrical division and full cytoplasm were selected and collected as 4-cell. After 72 hours, embryos with rounded morphology and full cytoplasm were selected and collected; since the time of cell development is not completely synchronized, embryos with from 8-cell to morula will be present at this stage. After culturing another 12 hours, the embryos with round shape and obvious cavity were selected and collected as blastocyst. So far, 6 stages including fertilized eggs, 2-cell, 4-cell, 8-16-cell, morula and blastocyst have all been collected.

#### 2.10 Immunofluorescence of mice embryos

The collected embryos at different stages were digested with 0.1% hyaluronidase (Beyotime Biotechnology, ST1384-25mg) at 37°C for 10 minutes. The digested embryos were fixed in 4% PFA (Beyotime Biotechnology, P0099-100ml) and kept overnight at 4°C. The fixed embryos were washed three times with 1 x PBS, then placed in 0.2% Triton-X-100, and infiltrated at room temperature for 30 minutes. After permeation, the fixed embryos were washed three times with 1 x PBS, and incubated in 1% bovine serum albumin (Amresco, Product No: 0332-100G) blocking solution for 30 minutes, and then moved to the primary antibody (RPL3, RPS3, NPM1, LSM14A or G3BP1) and incubate at 4°C overnight. After the embryos were incubated with a primary antibody, they were washed three times with 1 x PBS, and then incubated with a FITC-labeled Goat Anti-Rabbit IgG (H+L) (Beyotime Biotechnology, A0562) or abberior STAR RED (goat anti-mouse IgG: Item No. STRED-1001-20ug) for 2 hours and then washed three times with 1 x PBS, and stained with 5 mg/mL DAPI (Beyotime Biotechnology, C1005) for 15 minutes at room temperature. Wash three times with 1 x PBS, place the final processed embryos on a glass slide containing 5μL of mouting medium with antifading (Solarbio, Cat No. S2100), and analyze with a fluorescence microscope.

#### 2.11 OP-puromycin (OPP) Assay

NV stem cells were treated with either 1x PBS or 30μM OPP (APExBIO, A8778-5) for

30 min. Then the cells were washed with 1x PBS for three times and fixed with cold methanol for 2 min at -20 °C. The cells were washed with Tris buffered saline (TBS) and permeabilized with TBST (TBS with 0.2% Triton X-100) for 20 min at room temperature, and then washed with TBS again. Cells then were incubated with Alexa 448-azide (1:1000), 1 mM TCEP, 100μM TBTA, 1 mM CuSO4 for 1 h without light exposure, and washed with TBS for three times.

**2.12 Mitochondrial staining**

The mice embryos at different stages were digested with 0.1% hyaluronidase at 37°C for 15 minutes, and washed three times with 1x PBS, then incubated in M2 medium with Mito-tracker Red (100nM) (Thermal Fisher, M22425) for 15 min in a CO2 incubator at 37°C and 5% CO2. After incubation, the embryos were washed three times with 1x PBS, and imaged with 3D using super-resolution Zeiss LSM980.

Stem cells were directly incubated with Mito-tracker Green (100nM) (Thermal Fisher, M7514) for 15min in a CO2 incubator at 37℃ and 5% CO2. After incubation, the embryos were washed three times with 1x PBS, and imaged with 3D using super-resolution Zeiss LSM980.

**2.13 ER staining**

Stem cells were directly incubated with ER-Tracker (100nM) (Thermal Fisher, E12353) for 15 min in a CO2 incubator at 37°C and 5% CO2. After incubation, the embryos were washed three times with 1x PBS, and imaged with 3D using super-resolution Zeiss LSM980.

**2.14 FRAP Assay**

Fluorescence recovery after photobleaching (FRAP) is a method to qualitatively and quantitatively study biomolecule dynamics about movement of surrounding intact probes into the bleached spot in living cells. Cells were cultured in confocal dishes for two days, and stained by NucBlue™ Live ReadyProbes™ reagent (Thermal Fisher, R37605) for 15min, then washed three times with 1x PBS; at last, the live cells in live cell culture system were treated and imaged.

FRAP experiments were performed on a laser scanning confocal microscope (Nikon

A1) equipped with a 100× objective (N.A.=1.4). For living cells with expressing RPS18b-Zsgreen or RPL29-DsRed Ribo-macs were bleached using a 488 or 561 laser. A region of interest within the Ribo-macs was bleached using 15 mW laser irradiation for 1s, and the time-lapse images after photobleaching were collected for 300 s at a rate of 5s per frame. As for analysis of FRAP data, the fluorescent intensity (Itm) recorded on the bleached region in each time point (t) were normalized to fluorescent intensity (Itc) of nearby unbleached region, with the formula: It=(Itm/I0m)/(Itc/I0c ). Fluorescence recovery fraction for bleached intensity was further calculated with the formula: (It - Imin)/ (I0 - Imin). Imin is the unbleached fraction after photo-bleaching. Image J and Origin9 were used to measure and analyze the FRAP data.

#### 2.15 Teratoma Formation

5-10 million hiPSCs in a slurry of Matrigel and CDM medium (1:1) were injected into the subcutaneous of anesthetized NOD-SCID immunodeficient mice. Teratoma growth was monitored and teratoma size was made by quantifying approximate elliptical area (mm2) with the use of calipers measuring outward width and height.

#### 2.16 Teratoma Processing, Sectioning and HE (Hematoxylin and Eosin) staining

When the teratoma grew for about 70 days on average mice, the mice were euthanized by cervical dislocation. Tumor area was sprayed with 70% ethanol, and then teratoma was extracted via surgical excision using scissors and forceps. Teratoma was rinsed with 1x PBS, weighed, and photographed. Representative teratomas were cut into less than 5 mm diameter pieces and frozen in OCT for sectioning and HE staining.

Paraffin & frozen sectioning and HE staining was performed by Pathological section platform of the Institute of Biophysics, Chinese Academy of Sciences. In brief, teratomas after fixation for paraffin sectioning need gradient dehydration using ethanol (50%, 70%, 80%, 95%, 100%, 100%, 100%), and then were embed in fresh new paraffin and sectioned. Optimal Cutting Temperature (O.C.T.) blocks of teratomas for frozen sectioning were sectioned with a cryostat into 10micron sections onto a positively charged glass slide. The slide was then stained with HE staining. Sections from teratomas were confirmed to have the presence of all 3 germ layers: endoderm, mesoderm, and ectoderm via microscopy identification courtesy of pathologist Dr. Junfeng Hao.

Sections were fixed with 4% (w/v) paraformaldehyde for 15 min and permeabilized with 0.2% (v/v) Triton X-100 in PBS for 15 min. After blocking with 1% (v/v) BSA (Sangon Biotech, #9048-46-8) and 0.1% Tween 20（Sinopharm Chemical Reagent Co., Ltd., #30189328）for 60 min, cells were incubated with primary antibodies overnight at 4°C. Cells were washed with PBS for three times, then incubated with secondary antibodies and visualized by super-resolution microscopy including Zeiss LSM980, the images were taken at room temperature and images were analyzed using Imaris soft.

Figs. S1 to S5

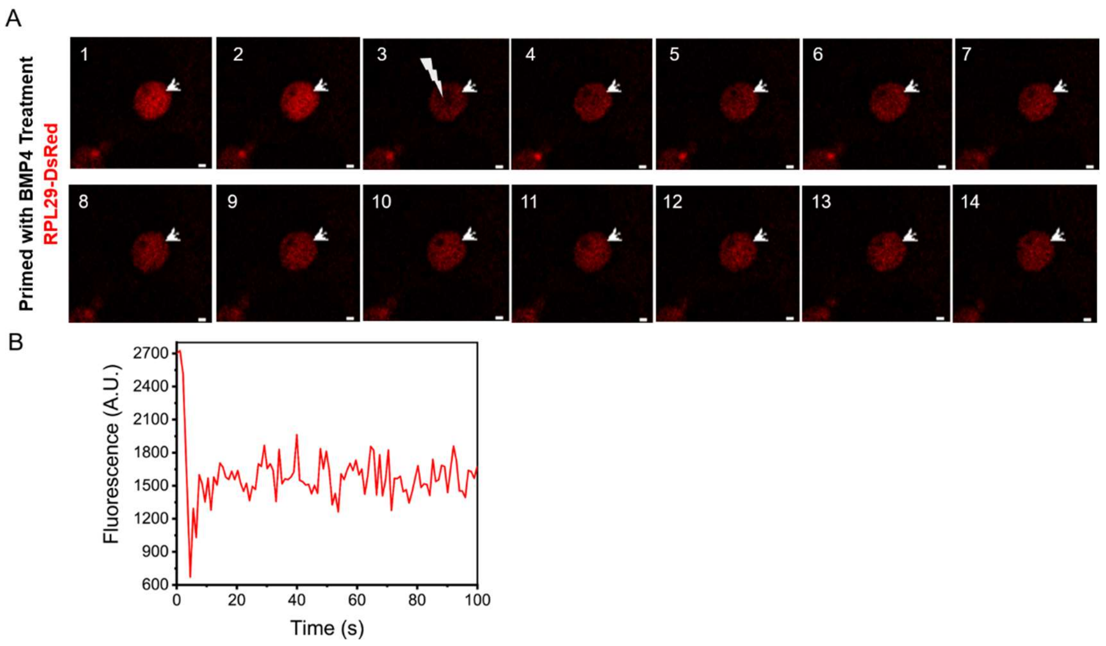


**fig. S1. FRAP assay in primed hiPSCs treated with BMP4.**

(A) and (B) Representative FRAP showing fluorescence of Ribo-macs could be recovered after photobleaching (white arrowheads, scale bar, 1um).

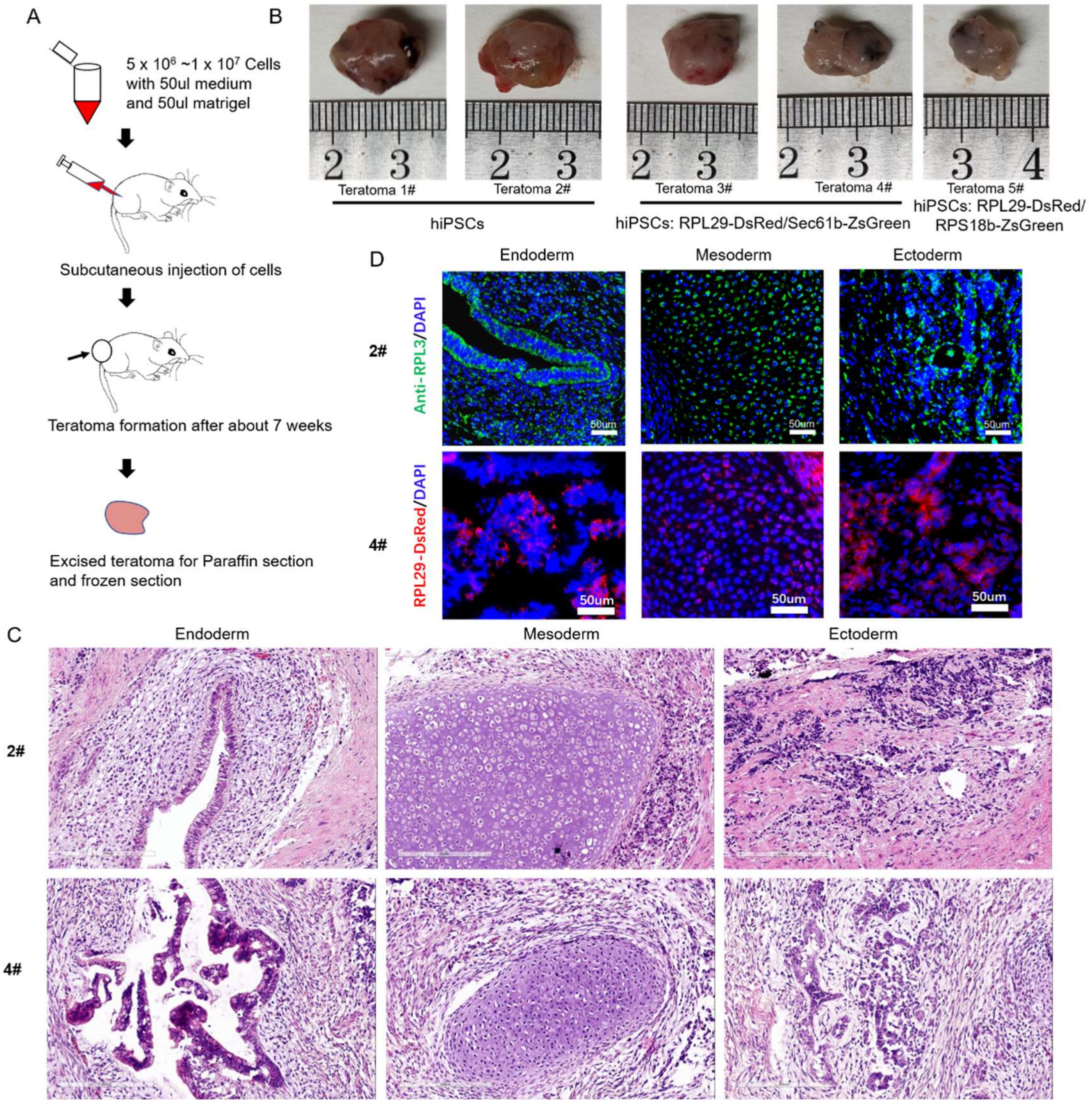


**fig. S2. Teratoma formation and characterization.**

(A) Schematic of general workflow. Subcutaneous injection of hiPSCs, hiPSCs with expressing RPL29-DsRed/Sec61b-ZsGreen and hiPSCs with expressing RPL29-DsRed/RPS18b-ZsGreen in a slurry of Matrigel and CDM medium was made in the right flank of NOD-SCID immunodeficient mice. Tumors were then extracted after about 7 weeks of growth. (B) Images of five teratomas generated. (C) H&E stains of the two teratoma histology sections. The presence of ectoderm, mesoderm, and endoderm confirmed for pluripotency and developmental potential. Scale bar, 200um. (D) Immunofluorescence staining for RPL3 and overexpression of RPL29-DsRed revealed that Ribo-macs exist in three germ layers, Scale bar, 50um.

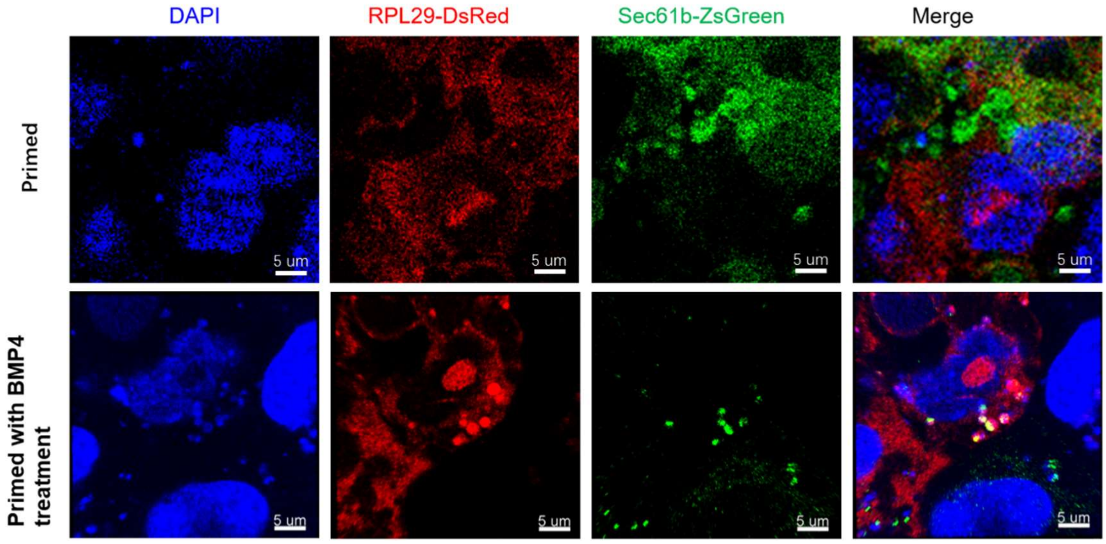


**fig. S3. Ribo-macs can produce proteins in primed hiPSCs treated with BMP4.**

Using expressing RPL29-DsRed and Sec61b-ZsGreen stem cells, the primed stem cells treated with BMP4 has Ribo-macs and Ribo-macs can synthesize proteins, which is identified by colocalization between RPL29-DsRed and Sec61b-ZsGreen, scale bar, 5um.

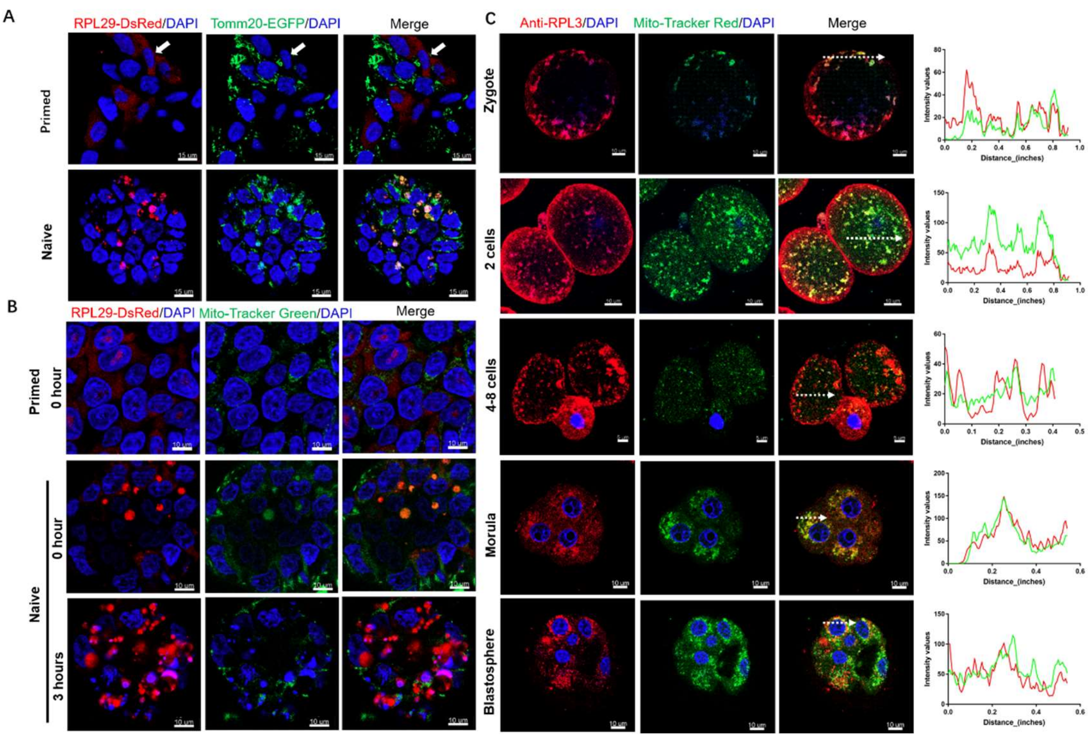


**fig. S4 Ribo-macs are related to mitochondrial biogenesis.**

(A) Observation of naïve stem cells with RPL29-DsRed/Tomm20-EGFP overexpression revealed colocalization of Ribo-macs and mitochondria but not in primed stem cells (white arrowheads), suggesting that Ribo-macs are related to mitochondria biogenesis. Scale bar, 15um. (B) Observation of colocalization of Ribo-macs and mitochondria at 0 hour between naïve and primed stem cells, and comparation of change of Ribo-macs and mitochondria at 0 hour and 3 hours in naive stem cells after mitochondrion staining using 100nM Mito-Tracker Green in expressing RPL29-DsRed stem cells. Scale bar, 10um. (C) To label mitochondria with Mito-Tracker red and immunofluorescence staining for RPL3 in five stages including zygote, 2 cells, 4-8 cells, morula and blastosphere of mouse embryos, and representative line profiles (dotted arrows) showed that colocalization of Ribo-macs and mitochondria, Scale bar, 10um or 5um.

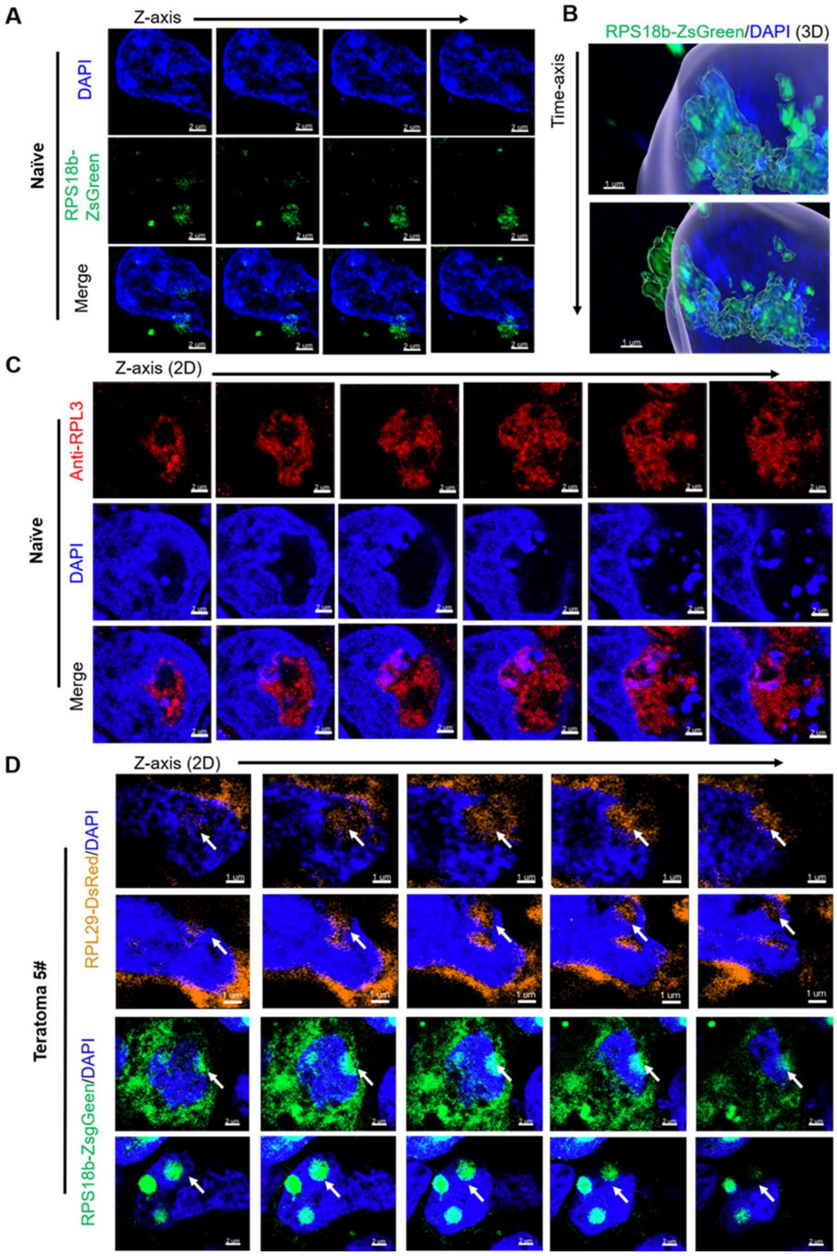


**Fig. S5. Nuclear export of ribo-macs is discovered in naïve stem cells and teratoma**

(A) Images of the second of (B) in the direction of Z-axis with 1um interval showing nuclear export of Ribo-macs (scale bar, 2um). (B) 3D images in the direction of time-axis with 5min interval showing nuclear export of Ribo-macs in naïve stem cells with expressing RPS18b-ZsGreen (scale bar, 1um). (C) Images of immunostaining for RPL3 in the direction of z-axis showing nuclear export of ribo-macs in naïve stem cells, (scale bar, 1um). (D) Representative images of RPS18b-ZsGreen (scale bar, 2um) and RPL29-DsRed (scale bar, 1um) in the direction of Z-axis with 1um interval showing nuclear export of Ribo-macs in teratoma (white arrowheads).

### Movies S1 to S21

**Movie S1.** 3D imaging of Ribo-macs in naïve stem cells based on immunostaining for RPS3.

**Movie S2.** 3D imaging of Ribosomes in primed stem cells based on immunostaining for RPS3.

**Movie S3.** 3D imaging of Ribo-macs in naïve stem cells based on immunostaining for RPS3 before conversion of naïve state to primed state.

**Movie S4.** 3D imaging of Ribosomes in primed stem cells based on immunostaining for RPS3 after conversion of naïve state to primed state.

**Movie S5.** 3D imaging of Ribo-macs in naïve stem cells with expressing RPL29-DsRed before conversion of naïve state to primed state.

**Movie S6.** 3D imaging of Ribosomes in primed stem cells with expressing RPL29-DsRed after conversion of naïve state to primed state.

**Movie S7.** FRAP imaging in naïve stem cells with expressing RPS18b-ZsGreen.

**Movie S8.** FRAP imaging in primed stem cells with expressing RPL29-DsRed treated by BMP4.

**Movie S9.** 3D imaging of Zygote based on immunostaining for RPS3.

**Movie S10.** 3D imaging of 2 cells based on immunostaining for RPS3.

**Movie S11.** 3D imaging of 4-8 cells based on immunostaining for RPS3.

**Movie S12.** 3D imaging of Morula based on immunostaining for RPS3

**Movie S13.** 3D imaging of Blastosphere based on immunostaining for RPS3.

**Movie S14.** 3D imaging of Ribo-macs in naïve stem cells with co-expressing RPL29-DsRed and RPS18b-ZsGreen.

**Movie S15.** 3D imaging of Ribo-macs in a naïve stem cell with co-expressing RPL29-DsRed and RPS18b-ZsGreen.

**Movie S16.** Live-cell imaging of Ribo-macs in a naïve stem cell with co-expressing RPL29-DsRed and RPS18b-ZsGreen.

**Movie S17.** Live-cell imaging of Ribo-macs in primed stem cells with co-expressing RPL29-DsRed and RPS18b-ZsGreen incubated with BMP4.

**Movie S18.** 3D imaging of Ribo-macs in naïve stem cells with expressing NPM1-mCherrry.

**Movie S19.** 3D imaging of Ribo-macs in primed stem cells with expressing NPM1-mCherrry.

**Movie S20.** 3D imaging of Ribo-macs in Blastosphere based on immunostaining for NPM1.

**Movie S21.** 3D imaging of Ribo-macs in Blastosphere based on immunostaining for NPM1.
